# Temperature and ecomorphology linked to blood pathogen incidence in neotropical amphibians

**DOI:** 10.64898/2026.07.07.736756

**Authors:** João Paulo de Oliveira Xavier, Diego Almeida-Silva, Arlei Marcili, Marcia Aparecida Sperança, Felipe Trovalim Jordão, Aline Diniz Cabral, Vanessa Kruth Verdade

## Abstract

While emerging diseases pose a global threat to amphibians, the dynamics of understudied vector-borne blood pathogens remain poorly understood. Pathogen occurrence is driven by a combination of environmental, ecological, and phylogenetic factors, yet how these drivers shape blood pathogen communities in tropical amphibians is largely unknown. In this study, we used molecular screening and phylogenetic linear models (PGLMMs) to evaluate how climate and ecomorphology influence the incidence of three blood pathogen groups (Trypanosomatidae, *Hepatozoon*, and *Rickettsia*) in wild anurans from a protected area in the Brazilian Atlantic Forest. Among 93 individuals sampled, over 93% were infected with at least one pathogen. *Trypanosomatidae* was the most common (76.3%), followed by *Rickettsia* (69.9%) and *Hepatozoon* (16.1%). Pathogen responses to temperature were contrasting: *Hepatozoon* occurrence increased in warmer periods, while *Trypanosomatidae* declined. Furthermore, rheophilic species showed a lower probability of *Rickettsia* infection, providing the first evidence that microhabitat use influences blood pathogen dynamics in amphibians. Our findings demonstrate that hemoparasite’s prevalence is driven by a multifaceted interplay of variables, highlighting that conservation strategies must account for these pathogen-specific responses to habitat use and environmental change, even within protected areas.

## INTRODUCTION

Amphibians are the most threatened terrestrial vertebrates (IUCN, 2025), facing global population declines from various stressors like habitat loss, pollution, climate change, and emergent diseases (Luedtke et al., 2023; Almeida-Silva et al., 2024). Their reliance on both aquatic and terrestrial habitats and permeable skin makes them vulnerable (Kiessecker, 2011; Awkerman et al., 2024). These factors increase mortality, reduce fitness, and inhibit population growth (Chambouvet et al., 2015; Sewell et al., 2021). Declining amphibian populations also impact ecosystems due to their roles as predators and prey (Zipkin & DiRenzo, 2022). Some of these threats have been extensively studied over the past few decades, especially focusing on frogs (Hof et al., 2011). However, the effects of anthropogenic stressors on anurans as hosts and reservoirs of enzootic and epizootic pathogens remain largely unknown.

A wide diversity of pathogens has been reported in anurans (Bower et al., 2019). The most studied are the fungus *Batrachochytrium dendrobatidis* and iridoviruses, directly implicated in amphibian declines (Piotrowski et al., 2004; Gray et al., 2009). These cases have been extensively analyzed, with research linking climate, land use, pollution, geography, and historical factors to host susceptibility (Lips et al., 2008; McCoy & Peralta, 2018; Santos et al., 2024). In contrast, hemoparasites remain comparatively overlooked, as their detection often requires histological or PCR analyses (Smith, 1996; Barrett et al., 2003). Hemoparasites comprise vector-borne blood parasites of diverse evolutionary origins, transmitted by leeches, dipterans, and ticks (Hamilton et al., 2007; Ferreira et al., 2015; Cotes-Perdomo et al., 2018), and are generally grouped into hemoflagellates, apicomplexans, and filarial nematodes (Davies & Johnston, 2000). Some hemoflagellates, such as trypanosomatids, are medically relevant because they also infect humans and other vertebrates (Barrett et al., 2003; Souza et al., 2014). Most trypanosomatids in anurans belong to the aquatic clade, a lineage associated with aquatic or semi-aquatic vertebrates (Ferreira et al., 2015; Isaak-Delgado et al., 2020). Among apicomplexans, *Hepatozoon* is notable for hosting the greatest diversity of anuran pathogens (Netherlands et al., 2018). Since it infects nearly all vertebrate groups (Smith, 1996), some species also hold zoonotic potential, affecting both domestic animals and wildlife.

Trypanosomatids and *Hepatozoon*, despite distinct taxonomy, share life cycle similarities. Both are transmissible via vectors and ingestion (Baneth et al., 2003; Roellig et al., 2010; Velásquez-Ortiz & Ramírez, 2021), with extracellular vector stages and intracellular vertebrate stages (Davies & Johnston, 2000; Baneth et al., 2003). Obligate intracellularity is also crucial for zoonotic pathogens like *Rickettsia*, common in frogs and reptiles (Sánchez-Montes et al., 2019). This parasitism originated early in *Rickettsia*’s evolution, aiding its diversification (Weinert et al., 2009). *Rickettsia* infect amphibians mainly through infected tick hematophagy (Cotes-Perdomo et al., 2018; Luz et al., 2018; Sánchez-Montes et al., 2019).

A set of environmental, phylogenetic, and ecological factors influence the occurrence and diversity of pathogens in vertebrate hosts (Arriero & Møller, 2008; Kamiya et al., 2014; Gutiérrez et al., 2019). Climate-related variables, for instance, play a key role in regulating both vector and amphibian host abundance due to temperature and precipitation fluctuations (Ficetola & Maiorano, 2016; Hernández-Ortiz et al., 2022). Warmer and wetter periods enhance amphibian breeding performance and population growth, resulting in more active hosts (Ficetola & Maiorano, 2016), while also increasing the abundance and activity of arthropod vectors in tropical regions (Legett et al., 2018; Hernández-Ortiz et al., 2022). In turn, evolutionary history can shape host susceptibility to blood pathogens, as phylogenetic constraints influence immune defenses and ecological interactions with vectors (Quillfeldt et al., 2011; Gupta et al., 2020). However, ecological factors, like differential susceptibility to parasites, have received comparatively little attention in studies on anurans. Amphibians exhibit distinct morphological adaptations associated with habitat use, allowing their classification into ecomorphs (Moen & Wiens, 2017). As environmental factors and stressors vary across microhabitats (Almeida-Silva et al., 2024), similar variation can be expected among parasite-vectors, and consequently different patterns of pathogen prevalence in different ecomorphs. Understanding the interplay of these factors is crucial for assessing and predicting changes in zoonotic pathogens affecting amphibians and other vertebrates, including humans.

The One Health concept offers a valuable framework that highlights the interconnection between human and environmental health by integrating principles from ecology, conservation, and public health (Destoumieux-Garzón et al., 2018). Amphibians, together with the broader aquatic community, play a central role in linking ecosystem components, contributing to functions such as food security and zoonotic disease regulation (Zipkin & DiRenzo, 2022). They are embedded in complex trophic networks that connect parasites, hosts, and vectors through predator–prey interactions (Harjoe et al., 2022). Although transmission pathways and their effects on host fitness remain poorly understood, blood pathogens have been associated with population declines and tadpole diseases worldwide (Davies & Johnston, 2000; Chambouvet et al., 2015). Efforts to a holistic perspective are therefore necessary to fully understand pathogen dynamics and to support the preservation of biodiversity, ecosystem integrity, and human health (Destoumieux-Garzón et al., 2018).

In this study, we examined the influence of environmental and ecomorphological factors on the prevalence of blood pathogens in anurans of a neotropical community, while also evaluating whether species-specific responses are independent or influenced by their evolutionary history using a phylogenetic generalized linear mixed model (PGLMM). By detecting three blood pathogen groups (Trypanosomatidae, *Hepatozoon*, and *Rickettsia*) through PCR assays we aimed to elucidate the relationship between parasite prevalence, climatic variation, and ecomorphological traits. We hypothesized that pathogen occurrence would be higher during warmer and wetter periods, driven by increased interactions between vectors and hosts. Furthermore, we predicted that ecomorphs would present different patterns of pathogen prevalence, and the rheophilic species would exhibit higher parasite probability due to their exposure to both aquatic and terrestrial vectors. Our findings provide new insights into the complex interplay between environmental factors, evolutionary constraints, and anuran-parasite dynamics, highlighting implications for conservation strategies under a rapidly changing climate.

## METHODS

### Study area

We conducted field work at Parque Natural Municipal Nascentes de Paranapiacaba (PNMNP), a protected area located in Santo André (São Paulo Metropolitan Region - Brazil; Figure 1). PNMNP has 426 hectares of regenerating secondary dense rainforest and is part of a bigger mosaic of protected areas, the Atlantic Forest Biosphere Reserve (RBMA, 2018). The region has one of the highest precipitation levels (about 3.300 mm annually) in the Atlantic Forest morphoclimatic domain, exhibiting two distinct climatic seasons: a warmer and wetter period (December–March) and a cooler, less humid period (April–November) (Gutjahr & Tavares, 2009).

**Figure 1:**
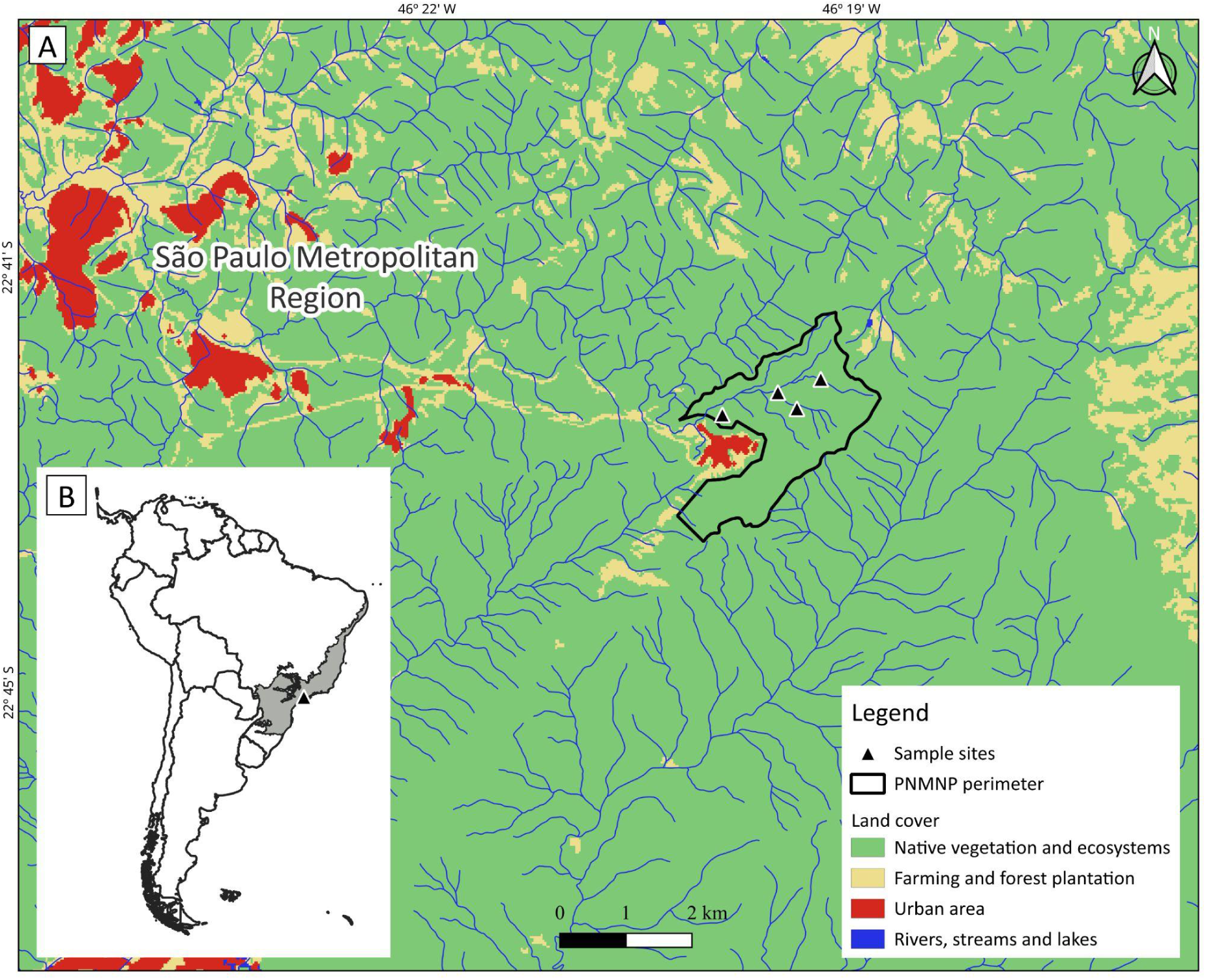
Study area. Location of frog sample sites at PNMNP (A), in Brazilian Atlantic Forest (gray), South America (B).

### Anuran sampling and categorization

We conducted our sampling over eight field campaigns from May to November 2015, with field trips every 20 days, totaling 32 days of fieldwork. Visits occurred during the driest months in the region, favoring frog sampling inside the streams and on its margins (Narvaes & Rodrigues, 2005). Transects (100m) were inspected in four streams in PNMNP, including 5m in the left and right terrestrial margins. Anurans were detected through visual and acoustic encounter surveys for two hours during the day and two hours at night. Distinct microhabitats were carefully scanned for anurans, such as rocks in streams, backwater, tree trunks, bromeliads, branches, and leaf litter (Doan, 2003). Voucher individuals were collected (collection permit SISBIO # 48099-1) and deposited at the zoological collection of Universidade Federal do ABC (ZUFABC). Frogs were identified to the species level (n = 16 species) following literature and specimen comparisons to voucher material at ZUFABC, and assigned to an ecomorphological group following the classification proposed by Haddad et al. (2013): Arboreal (n = 32, 7 spp.), Rheophilic (n = 15, 2 spp.), or Terrestrial (n = 46, 7 spp.). The diversity found represents approximately 20% of the anuran species richness registered for the region (Verdade et al., 2009; Trevine et al., 2014). Snout–vent length (SVL) was measured for each individual.

### Environmental variables

Climatic data were collected for the sampling period in the region. Precipitation data were obtained through the São Paulo Flood Alert System (SAISP), operated by Fundação Centro Tecnológico de Hidráulica (FCTH; SAISP, 2024). Additionally, temperature data were obtained from the Instituto Nacional de Meteorologia (INMET, 2024).

### Amphibian phylogenetic tree

Beyond microhabitat differences, phylogenetic relatedness may also influence species’ susceptibility to blood pathogens through parasite specialization (Ebert & Fields, 2020). To account for this, we inferred a phylogeny for our 19 species using 16S mitochondrial gene sequences (1,430 bp; SuppInfo2) from GenBank, prioritizing sequences from populations near the study area.

Sequences were aligned with MAFFT on the EMBL-EBI server (Li et al., 2015) using the Q-INS-i algorithm to account for secondary structure (Katoh & Toh, 2008). Model selection was performed with bModelTest under the transitionTransversionSplit set (Bouckaert & Drummond, 2017), and phylogenetic inference with divergence dating was implemented in BEAST v2.7.6 (Bouckaert et al., 2019). Two independent MCMC runs of 15 million generations each were performed, with a 25% burn-in.

Following Portik et al. (2023), divergence times were estimated under a strict molecular clock and Yule tree prior, with four fossil-calibrated nodes (Hyloidea: 66.0 Mya; Leptodactylidae: 40.4 Mya; Brachycephalidae: 34.0 Mya; Commutabirana: 5.4 Mya), modeled with lognormal priors reflecting fossil uncertainty. Clock rates were assigned weakly informative (1/x) priors.

MCMC convergence was verified with Tracer v1.7 (ESS > 200; Rambaut et al., 2018). After discarding burn-in, we summarized trees with TreeAnnotator v2.6.2 (Bouckaert et al., 2014) to produce a maximum clade credibility (MCC) tree with mean node heights. Both topology and divergence times were congruent with Portik et al. (2023).

### Blood pathogens detection

Liver samples from voucher specimens were stored at - 20 °C and used for parasite screening through molecular analyses. DNA was extracted with the DNeasy Blood & Tissue Kit (Qiagen, Hilden, Germany) following the manufacturer’s protocol, and purified using the GeneJET PCR Purification Kit (Thermo Fisher Scientific, Massachusetts, USA).

PCR reactions contained sterile water, forward and reverse primers (Table S1), dNTPs, and Taq DNA polymerase, with positive and negative controls. Each sample was tested separately with primers specific for Rickettsia (Labruna et al., 2004) and Hepatozoon (Almeida et al., 2012), while two primer sets were used for Trypanosomatidae detection (Hamilton et al., 2004; Jordão et al., 2021). Amplified products were visualized after agarose gel electrophoresis (150 V, 30 min) using a photo-documenter and digital camera.

### Statistical analyses

To investigate the relationship between the environmental drivers, ecomorphology, and blood pathogens occurrences in amphibians, we employed phylogenetic generalized linear mixed models (PGLMM). To account for potential evolutionary constraints, we incorporated a phylogenetic covariance matrix derived from the amphibian phylogeny as a random factor. To mitigate multicollinearity among predictors, we performed Pearson’s correlation analysis using the *cor* function in R (R Core Team, 2024), retaining only variables with coefficients <|0.7| (Figure S2; Dormann et al., 2013). Our analysis included three global models, each using the presence/absence of Trypanosomatidae, *Rickettsia*, or *Hepatozoon* as response variables. Continuous predictors included daily average precipitation, daily average temperature, and snout-vent length (SVL), while ecomorph categories served as categorical predictors. Sample sites (streams) were added as random factors to control for possible habitat quality variability. Models were fitted with binomial error distributions, suitable for binary response variables (Zuur et al., 2009).

AIC-based model selection guided the identification of the best-fit models, and phylogenetic mixed models were constructed using the *phyr* package (Li et al., 2020). To evaluate model performance, we calculated the proportion of variance explained by the models, considering both fixed and random effects (R^2^ and R^2^), using the *rr2* package (Ives, 2019). Model validation was performed with the *DHARMa* package (Hartig, 2016). All analyses were conducted in R version 4.4.0 (R Core Team, 2024).

## RESULTS

### Over 90% of amphibians presented blood pathogens

We conducted molecular analyses for blood pathogen detection in 93 individuals (16 species) out of 104 specimens (19 species) collected at the Parque Natural Municial Nascentes de Parnapiacaba (PNMNP; Table 1). From the 93 individuals, 93.6% (n = 87) tested positive for at least one of the three groups of parasites examined. Trypanosomatidae, *Hepatozoon* and *Rickettsia* were detected simultaneously in 8.6% (n = 8) of sampled frogs (Figure 3). Co-infections by Trypanosomatidae-*Rickettsia*, Trypanosomatidae-*Hepatozoon* and *Rickettsia-Hepatozoon* were found in 52.7% (n = 49), 11.8% (n = 11) and 12.9% (n = 12), respectively (Figure 3). Specifically, 76.3% (n = 71) of frogs were infected with trypanosomatids, 69.9% (n = 65) with *Rickettsia* and 16.1% (n = 15) with *Hepatozoon*. We found a detection frequency of 78.1% for trypanosomatids, 75.0% for *Rickettsia* and 15.6% for *Hepatozoon* among the arboreal ecomorphological group; a detection frequency of 66.7% for trypanosomatids, 53.3% for *Rickettsia*, and 20% for *Hepatozoon* among the rheophilic ecomorphological group; and 78.3% for trypanosomatids, 71.7% for *Rickettsia* and 15.2% for *Hepatozoon* among terrestrial ecomorphological group (Table 1).

**Table 1:** Prevalence (%) of blood pathogen infections (Trypanosomatidae, *Rickettsia*, and *Hepatozoon*) in anuran species. Data are categorized by host species and summarized by ecomorphological group (Arboreal, Rheophilic, and Terrestrial). *n* indicates the total number of individuals sampled in each category.

|  | n | Trypanosomatidae | *Rickettsia* | *Hepatozoon* | Total |
| --- | --- | --- | --- | --- | --- |
| **Arboreal** |  |  |  |  |  |
| *Aplastodiscus leucopygius* | 1 | 1 | 1 | 0 | 1 |
| *Bokermannohyla hylax* | 9 | 6 | 8 | 1 | 9 |
| *Dendrophryniscus imitator* | 7 | 6 | 6 | 0 | 7 |
| *Ischnocnema nigriventris* | 6 | 5 | 3 | 2 | 5 |
| *Ololygon rizibilis* | 1 | 1 | 1 | 1 | 1 |
| *Scinax alter* | 1 | 1 | 1 | 0 | 1 |
| *Vitreorana uranoscopa* | 7 | 5 | 4 | 1 | 7 |
| total | 32 | 25 | 24 | 5 | 31 |
| % |  | 78 | 75 | 16 | 97 |
| **Rheophilic** |  |  |  |  |  |
| *Hylodes asper* | 10 | 8 | 5 | 3 | 10 |
| *Hylodes caete* | 5 | 2 | 3 | 0 | 3 |
| total | 15 | 10 | 8 | 3 | 13 |
| % |  | 67 | 53 | 20 | 87 |
| **Terrestrial** |  |  |  |  |  |
| *Brachycephalus ibitinga* | 7 | 4 | 5 | 1 | 7 |
| *Haddadus binotatus* | 2 | 1 | 2 | 1 | 2 |
| *Ischnocnema aff. guentheri* | 10 | 8 | 7 | 0 | 10 |
| *Ischnocnema aff. parva* | 24 | 21 | 18 | 5 | 22 |
| *Physalaemus bokermanni* | 1 | 1 | 0 | 0 | 1 |
| *Physalaemus maculiventris* | 1 | 0 | 0 | 0 | 0 |
| *Rhinella icterica* | 1 | 1 | 1 | 0 | 1 |
| total | 46 | 36 | 33 | 7 | 43 |
| % |  | 78 | 72 | 15 | 93 |

### Temperature and ecomorphology influence on blood pathogens infections

The most plausible phylogenetic generalized linear mixed models (PGLMMs) indicate that the relationship between climatic drivers, ecomorphology, and infection occurrence varies among blood pathogens groups.

Trypanosomatidae infection probability decreased with higher daily average temperatures (β = −0.622, p = 0.0054; Figure 4, Table S2). Precipitation showed a marginally significant and weak negative association with Trypanosomatidae occurrence (β = −0.086, p = 0.0673; Table S2). Neither SVL nor ecomorphs had a significant effect on infection probability. No substantial phylogenetic signal (Var < 0.01, SD = 0.001; Figure 2, Table S2) or species-level variation (Var < 0.01, SD = 0.004; Table S2) was detected. However, some sampling sites exhibited slightly higher Trypanosomatidae infection rates than others (Var = 0.34, SD = 0.584; Table S2, Figure S1). Overall, random effects contributed minimally to the model’s explanatory power compared to fixed effects (R^2^ = 0.122, R^2^ = 0.158; Table S2).

**Figure 2:**
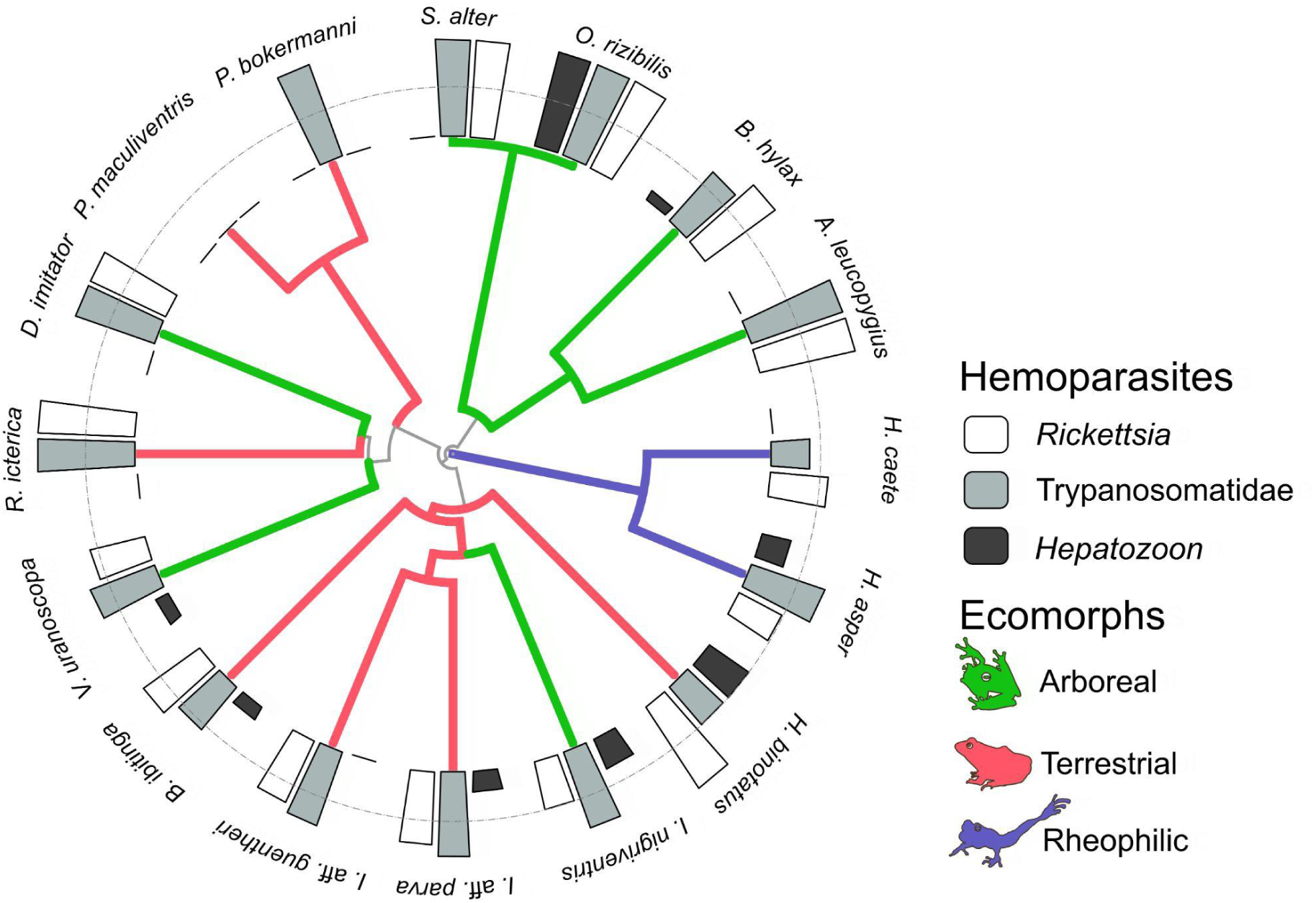
Phylogenetic tree of sampled anuran species showing the percentage of individuals infected by blood pathogens. Species are grouped by ecomorphs: arboreal (green), terrestrial (red), and rheophilic (blue).

**Figure 3:**
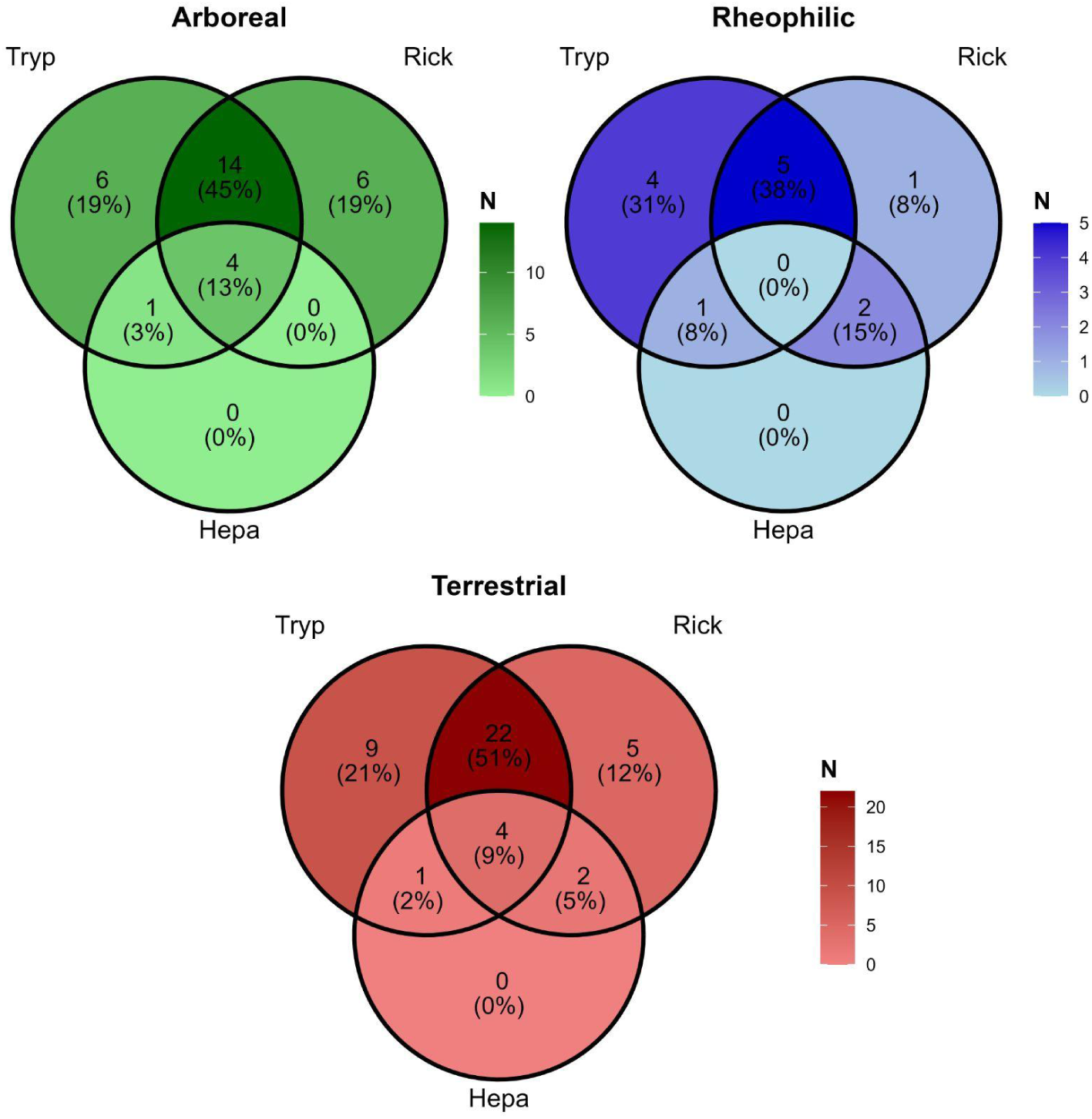
Co-infection patterns of Trypanosomatidae (Tryp), *Rickettsia* (Rick), and *Hepatozoon* (Hepa) in a neotropical amphibian assemblage, according to species habit: arboreal (green palette), rheophilic (blue palette), and terrestrial (red palette). Count and percentage of infected individuals are displayed inside the circles, with the overlaps indicating co-infections.

**Figure 4:**
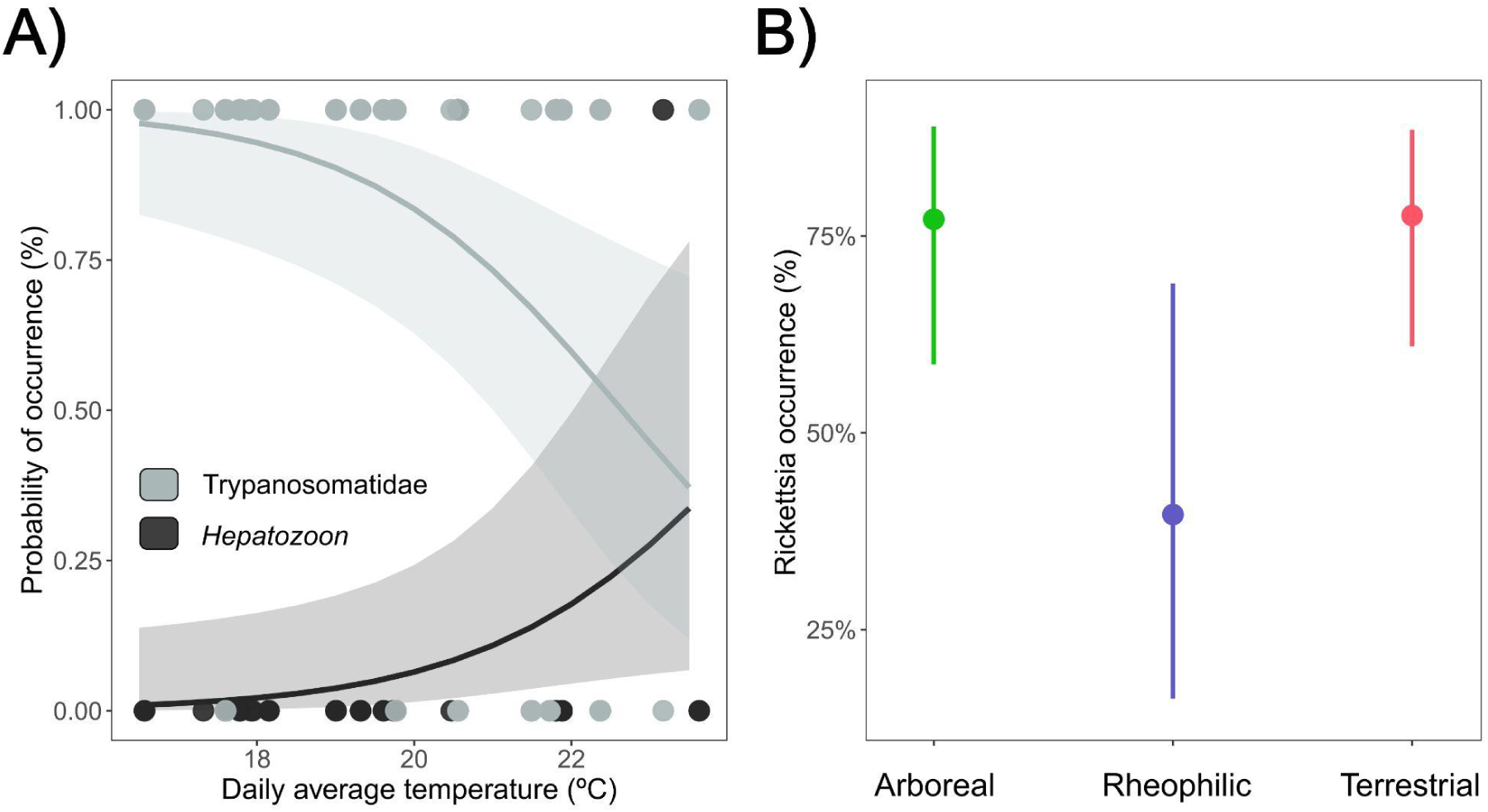
Relationship between the probability of blood pathogen infection with temperature and ecomorphologic groups. A) Probability of Trypanosomatidae (gray) and *Hepatozoon* (black) occurrence (%) as a function of daily average temperature (°C). B) Relationship between *Rickettsia* occurrence probability (%) and ecomorphs - arboreal (green), rheophilic (blue), and terrestrial (red).

The *Hepatozoon* infection probability increased with higher daily average temperatures (β = 0.552, *p* = 0.042; Figure 4, Table S3). Although not statistically significant, precipitation showed a weak positive association with *Hepatozoon* occurrence (β = 0.094, *p* = 0.076; Table S3). Neither SVL nor ecomorph categories had a significant effect on infection probability. Species-level variation had minimal influence on the model (Var < 0.01, SD = 0.003; Table S3), and a moderate variance may be explained by the phylogenetic random effect (Var = 0.10, SD = 0.315; Figure 2, Table S3). Sampling sites contributed substantially to the variation in infection probability (Var = 1.23, SD = 1.108; Table S3), with infections more concentrated in one of them (Figure S1). In general, random effects enhanced the model’s explanatory power (R^2^ = 0.128, R^2^ = 0.192; Table S3).

Rheophilic species had a significantly lower probability of *Rickettsia* infection than terrestrial and arboreal species (β = -1.636, *p* = 0.039; Figure 4, Table S4), which did not differ from each other. No meaningful associations were found between *Rickettsia* occurrence and SVL, precipitation, or temperature (Table S4). Random effects related to phylogenetic signal, species-level variation, and spatial distribution of sampling sites contributed minimally to the mode (Table S4)l. The explanatory power of the model increased only slightly with the inclusion of random effects and remained lower than that of the models for Trypanosomatidae and *Hepatozoon* occurrence (R^2^_lik_ = 0.078, R^2^_pred_ = 0.085; Table S4).

## DISCUSSION

Our work examined the influence of temperature, precipitation, and anuran ecomorphology on blood pathogen prevalence in a Neotropical amphibian community. Contrary to expectations, prevalence was not consistently higher during warmer and wetter periods, but instead followed more complex patterns: some infections were unrelated to temperature or precipitation (*Rickettsia*), while others showed opposing associations with temperature (Trypanosomatidae and *Hepatozoon*). Ecomorphology did not generally explain pathogen prevalence, except for the lower *Rickettsia* prevalence in rheophilic species, opposite to our initial hypothesis that stream-associated species would face higher exposure to both aquatic and terrestrial vectors.

The prevalence of trypanosomatid infections in our study area (∼76%) is comparable to rates reported in the Brazilian Pantanal (69%; Ferreira et al., 2007) and in an arboreal frog species from Argentina (73%; Pollo et al., 2023). Ferreira et al. (2007) reported 30% prevalence in Atlantic Forest sites with similar conditions to ours, with lower rates in drier areas (20%, Leal et al., 2009). Infection rates vary geographically: 20% in the Pantanal (Leal et al., 2009), and only 7% and 9% in Cerrado and Amazon frogs, respectively (Ferreira et al., 2015; Coelho et al., 2021). The high prevalence in PNMNP warrants further investigation.

Trypanosomatid infections are usually asymptomatic in amphibians but can cause histopathological lesions at high parasitemia levels (Sailasuta et al., 2011), and liver lesions are frequent in frogs from our study region (Almeida-Silva et al., 2024). Likely vectors include leeches and blood-feeding dipterans (Bartlett-Healy et al., 2009). Mosquitoes, sand flies, and frog-biting midges, known to transmit *Trypanosoma* and *Leishmania* to vertebrates, feed on amphibians (Borkent, 2008; Ferreira et al., 2008; Costa et al., 2021). The region’s high water availability (Gutjahr & Tavares, 2009) and landscape configuration likely support abundant dipteran populations (Marques et al., 2012). Assessing frog-biting insect diversity, abundance, and infection rates is thus a crucial subject to address at the locality.

*Rickettsia* prevalence in PNMNP amphibians (∼70%) was unexpectedly high. Since the early 2000s, infections have been reported in reptiles and amphibians worldwide (Sánchez-Montes et al., 2019), but mostly from ectoparasites, especially ticks, rather than directly in host tissues. Using ectoparasite infection as a proxy for host infection can be misleading: Horta et al. (2015) and Cotes-Perdomo et al. (2018) found *Rickettsia* DNA in 100% and 69.5% of ticks from toads, respectively, but none in the toads themselves. No ticks were found on frogs in PNMNP. While ticks are the main vectors, lice, mites, and fleas may also transmit *Rickettsia* (Kim, 2022). Its impact on amphibian fitness is unknown (Herczeg et al., 2021), but there’s evidence that reptiles can act as reservoirs for zoonotic species (Whitworth et al., 2003). Clarifying vector-host relationships and health effects requires further study.

*Hepatozoon* prevalence (16.13%) falls within the range reported for other Brazilian regions: 6.06% in *Leptodactylus* species from the Cerrado biome (Úngari et al., 2021), and 28.96% in Pantanal anurans (Leal et al., 2015). Flies, mosquitos, ticks, mites, and leeches, are known to transmit *Hepatozoon* to amphibians and reptiles (Smith, 1996; Leal et al., 2015; Úngari et al., 2021) and infections have been linked to inflammatory lesions in frogs (Sailasuta et al., 2011). Almeida-Silva et al. (2024) reported liver lesions in PNMNP frogs consistent with inflammatory responses, potentially linked to toxic elements in local waters. Thus, comprehending the infection cycle and its fitness implications is essential.

This is the first study to jointly assess these three parasite groups in amphibians. Total prevalence (93%) exceeds reports from elsewhere in South America, where the highest rate is ∼70% (Pollo et al., 2023). Co-infection with all three pathogens occurred in 8.6% of individuals - its significance for Neotropical communities remains unclear. As in our findings, trypanosomatid infections (26%) exceeded apicomplexan hemoparasites (7%) in eastern Amazon frogs (Pinho et al., 2021). We found mixed trypanosomatid-apicomplexan infections in 11.8% of frogs, nearly twice the 6.2% reported by Pinho et al. (2021). *Hepatozoon*-*Rickettsia* co-infections (12.9%) were slightly more common than *Hepatozoon-*Trypanosomatidae, perhaps due to their opposite temperature associations.

Research on amphibians and other vertebrates suggests a positive association between hemoparasites prevalence and temperature (Davies & Johnston, 2000; Bartlett-Healy et al., 2009; Sehgal, 2015). Increases in temperature are known to enhance vector metabolism, feeding frequency, and population growth (Legett et al., 2017; Hernández-Ortiz et al., 2022), all of which can elevate transmission rates. Moreover, amphibians are generally more active during warmer periods and tend to remain near insect oviposition sites, further increasing their exposure to vectors (Wood et al., 2007). Experimental data indicate that the reproduction of *Hepatozoon clamatae*, a species infecting frogs, is favored by warmer temperatures (Trites et al., 2013). Climate models predict rising temperatures in the region (Masson-Delmotte et al., 2021), which could increase infections and zoonotic potential under a One Health perspective (Short et al., 2017).

Despite these patterns, our results show complexity: Hepatozoon prevalence rose with temperature, Rickettsia showed no temperature relationship, and Trypanosomatidae prevalence decreased. Limited data on vector communities, transmission routes (e.g., ingestion vs. hematophagy), and parasite dynamics constrain interpretations. In parallel, temperature-driven parasite migration to peripheral vessels (Bardsley & Harmsen, 1969) could affect detectability depending on the point of blood sampling on frogs. Surveys of vectors and pathogen prevalence in a broader geographic scope are urgently needed to clarify the ecological drivers underlying these patterns.

Aside from *Rickettsia*’s lower prevalence in rheophilic species, ecomorphology had no clear effect on Trypanosomatidae or *Hepatozoon*. Considering ticks are the primary transmitters of rickettsial diseases, tick ecology and distribution may explain lower *Rickettsia* in stream-dwellers, with terrestrial/arboreal species more exposed (Andoh et al., 2015; Sánchez-Montes et al., 2019). Rheophilic amphibians, however, are more prone to waterborne parasites (Kriger & Hero, 2007), highlighting habitat’s role in some pathogen groups.

We did not find strong phylogenetic signals in our data, but only a moderate variation in *Hepatozoon* prevalence. Phylogenetic signals highlight the influence of conserved morphological, physiological, biochemical, and ecological traits shared among closely related species on parasite dynamics (Huang et al., 2014). Indeed, host evolutionary history influences hemoparasite diversity in birds and mammals (Huang et al., 2014; Clark et al., 2018). Identifying the specific host characteristics that drive parasite occurrence, beyond those examined here, will require further interdisciplinary research incorporating a broader range of traits. Our limited phylogenetic scope (16 species) likely constrained signal detection; larger-scale studies may reveal new patterns.

Finally, the PNMNP is a touristic area near the São Paulo Metropolitan Region (Figure 1) with considerable proximity among people, domestic animals, and wildlife, and is a well-known locality from its anurofauna perspective (Verdade et al., 2009). The region has a history of environmental pollution from a nearby industrial complex (Lemos, 1998), and despite control measures, high contaminant levels persist (CETESB, 2023), which have been recently associated with histopathological lesions in local anurans (Almeida-Silva et al., 2024). Compromised immune responses (Linzey et al., 2003) linked to chemical pollutants could explain the high pathogen prevalence registered in our study; on the other hand, hemoparasites are also vulnerable to waterborne contaminants that can cause cellular damage (Silva et al., 2005). Although geographic variation in pollutant exposure was not measured, the studied blood pathogens belong to groups with close relatives that infect other vertebrates, including humans, and their zoonotic potential remains unevaluated.

## CONCLUSION

Our study reveals a high prevalence (>93%) and complex occurrence patterns of understudied blood pathogens in a neotropical amphibian community, underscoring that even protected areas are not free from disease risks. Pathogen responses to environmental and ecological drivers proved highly context-dependent: temperature showed opposite associations with *Hepatozoon* and Trypanosomatidae incidence, while host ecomorphology was linked to lower *Rickettsia* infection risk in rheophilic anurans. Such complexity challenges simplistic predictions about climate change effects on wildlife diseases and suggests that factors shaping susceptibility to well-studied pathogens may not apply to hemoparasites. Critical gaps remain regarding vector identity, host fitness impacts, and potential zoonotic risks, especially as global change reshapes disease dynamics. Addressing these questions is essential for developing effective conservation strategies that incorporate host-specific susceptibility and for advancing our understanding of tropical ecosystem health under growing anthropogenic pressures.

## Supporting information

SuppInfo1

SuppInfo2

## Acknowledgements

This work was supported by Conselho Nacional de Desenvolvimento Científico e Tecnológico (CNPq), Coordenação de Aperfeiçoamento de Pessoal de Nível Superior (CAPES), and Fundação Universidade Federal do ABC (UFABC). We would like to thank ICMBIO for the collection permit (SISBIO #48099-1). Our gratitude also extends to the Municipality of Santo André and the management of Parque Natural Municipal Nascentes de Paranapiacaba (PNMNP) for approving the project (administrative process 47256/2014-3). Special thanks to Ingo Grantsau and Leandro Wada Simone, both from the PNMNP.

## Author contributions

João Paulo de Oliveira Xavier, Diego Almeida-Silva and Vanessa Kruth Verdade conceptualised the study; Vanessa Kruth Verdade, Márcia Aparecida Sperança and Arlei Marcili provided logistics, funding and institutional support; Diego Almeida-Silva and João Paulo de Oliveira Xavier conducted field work and frog sampling; Márcia Aparecida Sperança, João Paulo de Oliveira Xavier, Arlei Marcili, Felipe Trovalim Jordão and Aline Diniz Cabral performed molecular analyses; Diego Almeida-Silva built the amphibian phylogenetic tree; João Paulo de Oliveira Xavier analysed the data; João Paulo de Oliveira Xavier led the writing of the manuscript, with all authors contributing critically to the drafts, notably Vanessa Kruth Verdade and Diego Almeida-Silva. All authors gave final approval for publication.

## Data availability statement

Data and code used in the study are available at *Zenodo* (Xavier et al., 2025; https://doi.org/10.5281/zenodo.16763574).

