## Supplementary material for "Temperature and ecomorphology linked to blood pathogen incidence in neotropical amphibians": SuppInfo1

**Supporting Information**

**Table S1:** Description of the oligonucleotides used in the molecular analyses.

|  |  |  |  |
| --- | --- | --- | --- |
| **Hemoparasite** | **Gene name** | **5’-3’** | **Reference** |
| *Rickettsia* | CS-62F | GCAAGTATCGGTGAGGATGTAAT | Labruna et al.,  2004 |
|  | CS-462R | GCTTCCTTAAAATTCAATAAATCAGGAT | Labruna et al.,  2004 |
| *Hepatozoon* | HEP1 | CGCGAAATTACCCAATTCA | Almeida et al.,  2012 |
|  | HEP4 | TAAGGTGCTGAAGGAGTCGTTTAT | Almeida et al.,  2012 |
| Trypanosomatidae | Quit224F | GTTCCACTACGAGGCCTTC  TTCAA | Jordão et al., 2021 |
|  | Quit1182R | CAGATCATTATCCCAGACAAGTT | Jordão et al., 2021 |
|  | GAP3F | GTGAAGGCGCAGCGCAAC | Hamilton et al., 2004 |
|  | GAP5R | CCGAGGATGYCCTTCATG | Hamilton et al., 2004 |

**Table S2:** Parameters and metrics of the most plausible Phylogenetic Generalized Linear Mixed Model (PGLMMs) for Trypanosomatidae occurrence data, selected based on AIC. 1|sp = Random effect for species; 1|sp__ = Random effect for phylogenetic relationships; 1|stream = sample site random effect; habitRheophilic = Rheophilic ecomorph; habitTerrestrial = Terrestrial ecomorph; SVL = snout-vent length; precip = Daily average precipitation; mean = Daily average temperature; SE = Standard Error; SD = Standard deviation; Exp. Var. = Explained variance; R^2^lik = Variance of fixed components; R^2^pred = Variance of fixed and random components; . = Marginally significant; * = Statistically significant; ** = Highly significant.

|  |  |  |  |  |
| --- | --- | --- | --- | --- |
| **Trypanosomatidae** |  |  |  |  |
| Random effects: |  |  |  |  |
|  | **Variance** | **SD** |  |  |
| 1\|sp | 1.600e-05 | 0.004 |  |  |
| 1\|sp__ | 2.259e-06 | 0.001 |  |  |
| 1\|stream | 3.406e-01 | 0.584 |  |  |
| Fixed effects: |  |  |  |  |
| **Parameter** | **Estimate (β)** | **SE** | **z value** | ***p* value** |
| Intercept | 14.787992 | 4.966770 | 2.9774 | 0.002907 ** |
| habitRheophilic | -0.084954 | 0.800470 | -0.1061 | 0.915479 |
| habitTerrestrial | -0.121239 | 0.635387 | -0.1908 | 0.848674 |
| SVL | -0.014482 | 0.016290 | -0.8890 | 0.374002 |
| precip | -0.085174 | 0.046543 | -1.8300 | 0.067250 . |
| tmean | -0.621763 | 0.223732 | -2.7790 | 0.005452 ** |
| Performance evaluation: |  |  |  |  |
| **R²** | **Exp. Var.** |  |  |  |
| R²lik | 0.1216 |  |  |  |
| R²pred | 0.1577 |  |  |  |

**Table S3:** Parameters and metrics of the most plausible Phylogenetic Generalized Linear Mixed Model (PGLMMs) for *Hepatozoon* occurrence data, selected based on AIC. 1|sp = Random effect for species; 1|sp__ = Random effect for phylogenetic relationships; 1|stream = sample site random effect; habitRheophilic = Rheophilic ecomorph; habitTerrestrial = Terrestrial ecomorph; SVL = snout-vent length; precip = Daily average precipitation; mean = Daily average temperature; SE = Standard Error; SD = Standard deviation; Exp. Var. = Explained variance; R^2^lik = Variance of fixed components; R^2^pred = Variance of fixed and random components; . = Marginally significant; * = Statistically significant; ** = Highly significant.

| ***Hepatozoon*** |  |  |  |  |
| --- | --- | --- | --- | --- |
| Random effects: |  |  |  |  |
|  | **Variance** | **SD** |  |  |
| 1\|sp | 9.463e-06 | 0.003076 |  |  |
| 1\|sp__ | 9.950e-02 | 0.315435 |  |  |
| 1\|stream | 1.229e+00 | 1.108389 |  |  |
| Fixed effects: |  |  |  |  |
| **Parameter** | **Estimate (β)** | **SE** | **z value** | ***p* value** |
| Intercept | -14.2658405 | 6.0103551 | -2.3735 | 0.01762 * |
| habitRheophilic | -0.1603814 | 1.0923072 | -0.1468 | 0.88327 |
| habitTerrestrial | 0.4641409 | 0.7988043 | 0.5810 | 0.56121 |
| SVL | 0.0038717 | 0.0255065 | 0.1518 | 0.87935 |
| precip | 0.0942637 | 0.0531987 | 1.7719 | 0.07641 . |
| tmean | 0.5519989 | 0.2707984 | 2.0384 | 0.04151 * |
| Performance evaluation: |  |  |  |  |
| **R²** | **Exp. Var.** |  |  |  |
| R²lik | 0.1283 |  |  |  |
| R²pred | 0.1920 |  |  |  |

**Table S4:** Parameters and metrics of the most plausible Phylogenetic Generalized Linear Mixed Model (PGLMMs) for *Rickettsia* occurrence data, selected based on AIC. 1|sp = Random effect for species; 1|sp__ = Random effect for phylogenetic relationships; 1|stream = sample site random effect; habitRheophilic = Rheophilic ecomorph; habitTerrestrial = Terrestrial ecomorph; SVL = snout-vent length; precip = Daily average precipitation; mean = Daily average temperature; SE = Standard Error; SD = Standard deviation; Exp. Var. = Explained variance; R^2^lik = Variance of fixed components; R^2^pred = Variance of fixed and random components; . = Marginally significant; * = Statistically significant; ** = Highly significant.

| ***Rickettsia*** |  |  |  |  |
| --- | --- | --- | --- | --- |
| Random effects: |  |  |  |  |
|  | **Variance** | **SD** |  |  |
| 1\|sp | 4.405e-06 | 0.0020988 |  |  |
| 1\|sp__ | 0.000e+00 | 0.0000000 |  |  |
| 1\|stream | 8.732e-07 | 0.0009344 |  |  |
| Fixed effects: |  |  |  |  |
| **Parameter** | **Estimate (β)** | **SE** | **z value** | ***p* value** |
| Intercept | -1.160136 | 3.263111 | -0.3555 | 0.72219 |
| habitRheophilic | -1.636433 | 0.796546 | -2.0544 | 0.03994 * |
| habitTerrestrial | 0.027392 | 0.565994 | 0.0484 | 0.96140 |
| SVL | 0.035705 | 0.025164 | 1.4189 | 0.15593 |
| precip | 0.057009 | 0.043484 | 1.3110 | 0.18984 |
| tmean | 0.056164 | 0.148660 | 0.3778 | 0.70558 |
| Performance evaluation: |  |  |  |  |
| **R²** | **Exp. Var.** |  |  |  |
| R²lik | 0.0780 |  |  |  |
| R²pred | 0.0849 |  |  |  |

**Figure S1:** Distribution of infected individuals in the four streams sampled at PNMNP. Tryp = Positives for Trypanosomatidae; Hepa = Positives for Hepatozoon; Rick = Positives for Rickettsia; UninInd = Uninfected individuals.

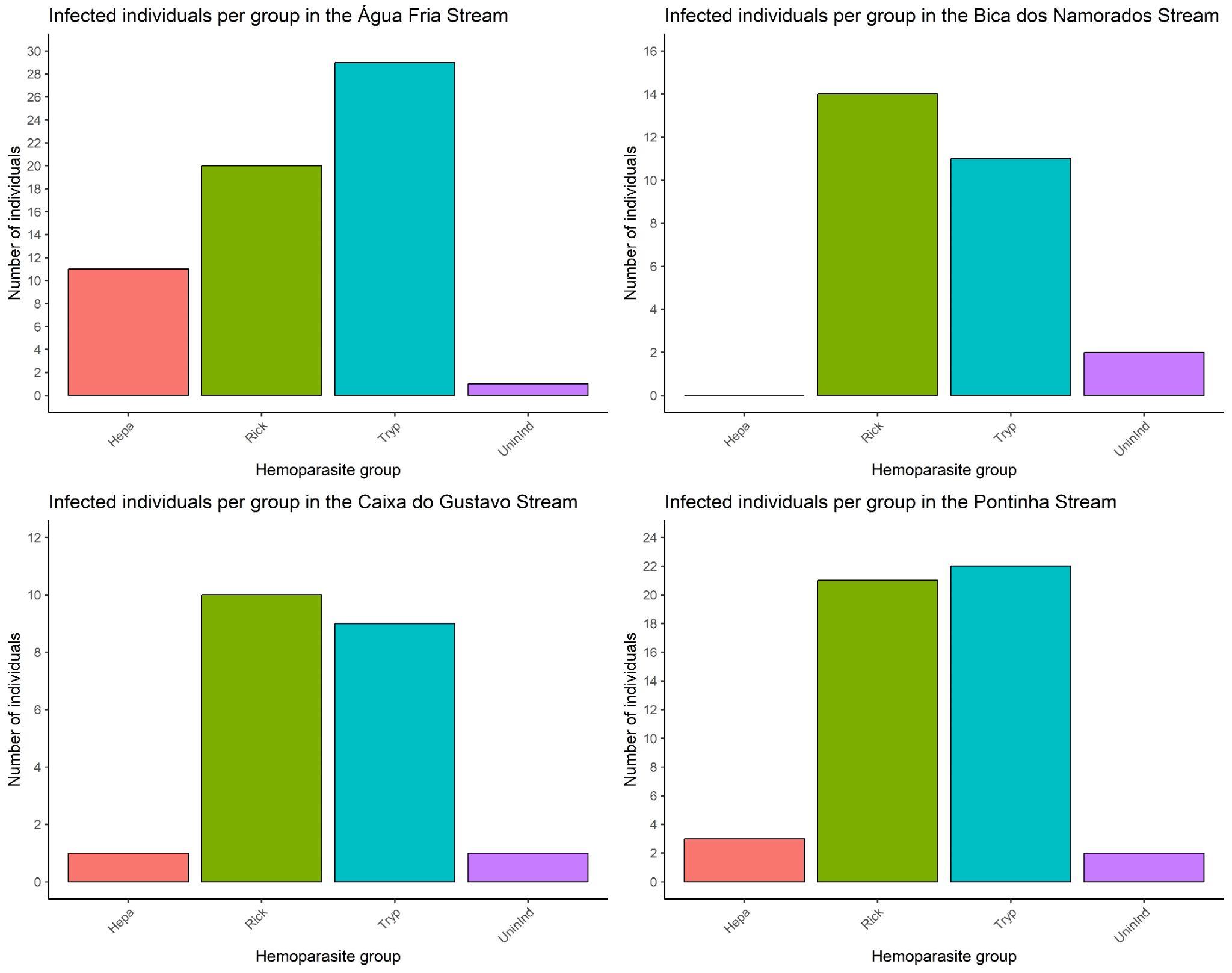

**Figure S2:** Correlation between the predictor variables used in the models. Only variables with a correlation lower than |0.7| were retained. Precip = Daily average precipitation; tmean = Daily average temperature; SVL = Snout-vent length; Mass = Body mass. The chart was made using the *‘corrplot’* function (Wei & Simko, 2024) in RStudio (R Core Team, 2024).

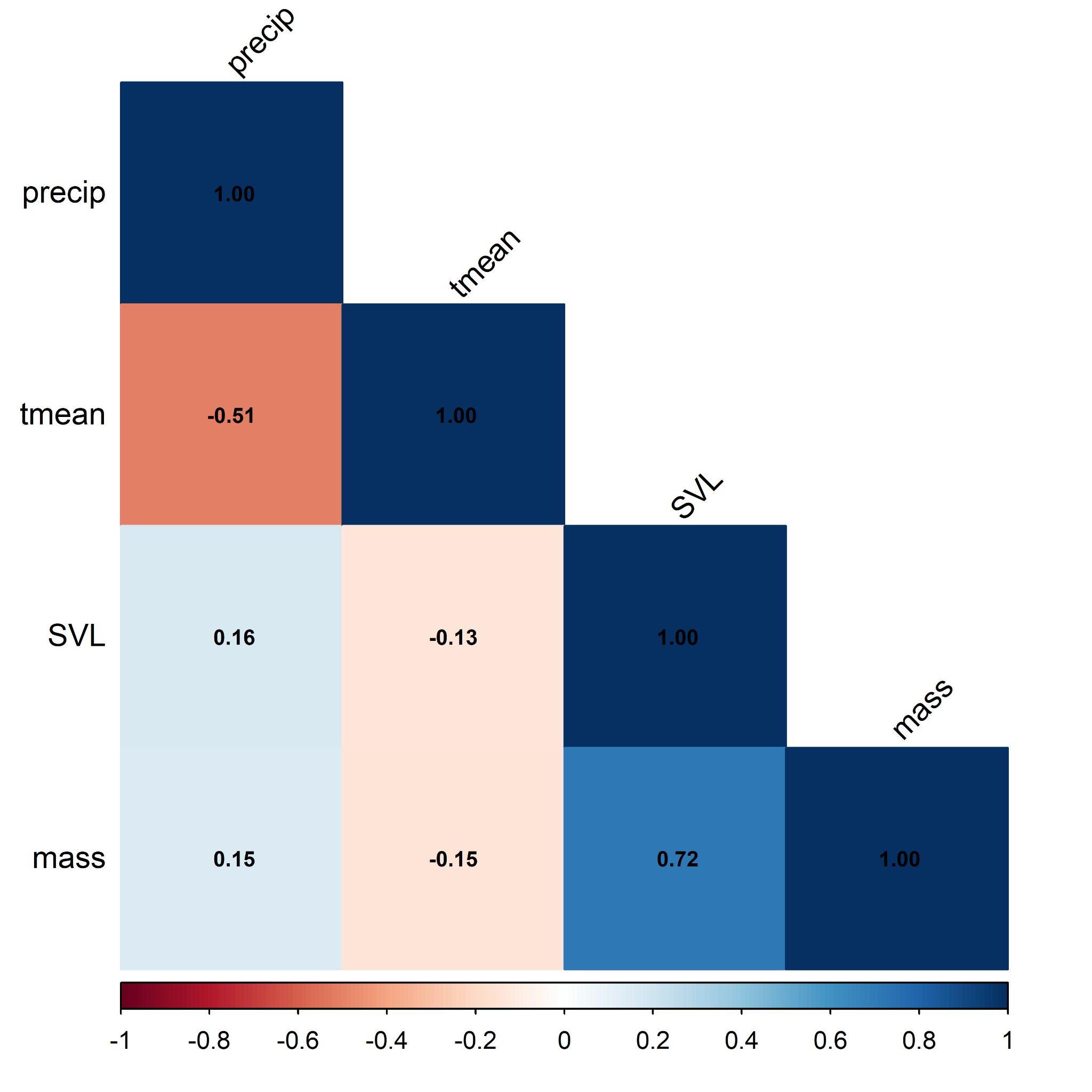
